# *In silico* evidence suggests that the SARS-CoV-2 Spike protein may target coiled-coil regions of numerous cytoskeletal and cytoskeleton-associated proteins

**DOI:** 10.1101/2025.01.20.633897

**Authors:** Pierre Brézellec

## Abstract

Understanding the interactions between host and viral envelope proteins is essential to get insights into the dynamics of viral infection. To investigate more closely the mechanisms governing SARS-CoV-2 entry and intracellular trafficking, I reanalyzed the most extensive SARS-CoV-2–human protein–protein interactome dataset currently available. My investigation centered on the Spike S protein, a key player in initiating viral infection by binding to the host cell membrane receptor Angiotensin-Converting Enzyme 2 (ACE2). I first present evidence demonstrating the statistical overrepresentation of actin-binding proteins among the Spike S partners/interactors. Next, I show that a majority of these partners contains a structural domain sharing high similarity with the C-terminal region of the Myosin II heavy chain, Myosin II being known for its roles in muscle contraction and various cellular motility processes. I subsequently demonstrate that this domain is particularly prevalent in actin-binding proteins, intermediate filaments proteins and kinesins, which all are related to the cytoskeleton known to be involved in diverse cellular functions, including endocytosis and intracellular transport — processes crucial for viral infections. Finally, I highlight that the structural domain mentioned above is a bonafide coiled-coil region. I therefore conclude that Spike S might target proteins possessing such regions. Collectively, my findings suggest that the interactions between SARS-CoV-2 Spike S and human proteins, potentially mediated by coiled-coil regions, may have been underestimated. As this work relies on *in silico* evidence, direct biological extrapolations require caution.

## Introduction

Exploring protein-protein interactions (PPI) between host proteins and viral proteins has emerged as a relevant approach in viral infection research (Calderwood *et al*., 2007). In the context of COVID-19, such approaches have been used to identify pharmacological agents with antiviral activity or potential host therapeutic targets (Gordon *et al*., 2020, Zhou *et al*., 2022). In this study, I focused on the most comprehensive and reliable SARS-CoV-2–human PPI data available to date (Zhou *et al*., 2022), based on both two-hybrid (Y2H) and affinity purification-mass spectrometry (AP-MS), to enhance our understanding of virus infection dynamics (encompassing entry and propagation both intra-and extracellularly).

Large-scale protein interaction studies using techniques like Y2H (Rajagopala *et al*., 2015) and AP-MS (Causier *et al*., 2004) have well-documented limitations. For example, they might miss transient interactions or detect interactions between proteins that are not normally found in the same cellular compartment or not expressed at the same time in the cell, *etc*. (<u>EBI Training Online Courses</u>, (Millan, 2022)). Despite these weaknesses, however, Y2H and AP-MS often generate high-quality data which can be used as a reliable and robust basis for further investigation. In our context, at first glance, this assertion might seem surprising given a well-known major limitation of the aforementioned interactome: the critical interaction between the Spike S binding domain (RBD) and the host Angiotensin-Converting Enzyme 2 (ACE2), which allows for the initiation of viral infection, has not been identified. However, while this latter interaction is clearly of great importance, this interactome can potentially help to detect or reveal other types of interesting and meaningful interactions, such as those involving other regions of Spike S, or those taking place at other stages of the infection process.

Relying on the Spike S sub-network of the aforementioned interactome, I first demonstrate that actin-binding proteins are statistically overrepresented within the set of Spike S interactors/partners. Using Phyre2 (Kelley *et al*., 2015), a popular bioinformatics method for protein structure prediction, I then show that a majority of Spike S partners share structural similarities with the C-terminal part of the Myosin II heavy chain, which is a well-documented coiled-coil region (Rahmani *et al*., 2021). Next, using BackPhyre, a tool that identifies proteins harboring a given structural domain, I identify a set of human proteins containing with high confidence the C-terminal part of the Myosin II heavy chain and demonstrate that this set is enriched in cytoskeletal and associated cytoskeletal proteins, with actin-binding proteins, intermediate filament proteins, and kinesins all being statistically overrepresented compared to their distribution in the human proteome (refer to the Figure for an overall view of my results).

My work thus provides *in silico* evidence suggesting that SARS-CoV-2 Spike S might target proteins harboring coiled-coil regions, which, as I show here (see also (Simm *et al*., 2021)), are particularly abundant in cytoskeletal and cytoskeletal-associated proteins. This suggestion is in line with numerous studies that have highlighted the crucial role of the host cell cytoskeleton for coronavirus infection, including SARS-CoV-2 (Wen *et al*., 2020; Kloc *et al*., 2022). For example, the actin-binding protein Myosin 9 (Chen *et al*., 2021) and the intermediate filament vimentin (Amraei *et al*., 2022) have been shown to facilitate SARS-CoV-2 host cell entry.

## Materials and methods

### Interactome data

The interactions between the Spike S protein and human proteins used in this paper are extracted from a protein-protein interactome published in Nature Biotechnology (Zhou *et al*., 2022). They involved 22 human proteins, hereafter denoted as interactors or partners, as shown in Table 1 below.

**Table 1.** Each line represents a Spike S partner. The ‘UniProtID’ column identifies the protein using its unique identifier in the UniProt database. The ‘Protein Name’ column indicates the name of the protein. The ‘Technique of identification’ column refers to the approach used to identify the considered interaction, where ‘Y2H’ denotes/stands for ‘Yeast Two-Hybrid’ and ‘AP-MS’ stands for ‘Affinity Purification-Mass Spectrometry’.

| UniProt ID | Protein Name | Technique of identification |
| --- | --- | --- |
| ACTN4_HUMAN | Alpha-actinin-4 | Y2H |
| MYH9_HUMAN | Myosin-9 | AP-MS |
| MYH10_HUMAN | Myosin-10 | AP-MS |
| MYH14_HUMAN | Myosin-14 | AP-MS |
| TPM1_HUMAN | Tropomyosin alpha-1 chain | AP-MS |
| TPM2_HUMAN | Tropomyosin beta chain | AP-MS |
| TPM3_HUMAN | Tropomyosin alpha-3 chain | AP-MS |
| TPM4_HUMAN | Tropomyosin alpha-4 | AP-MS |
| COR1C_HUMAN | Coronin-1C | AP-MS |
| TMOD3_HUMAN | Tropomodulin-3 | AP-MS |
| BICL2_HUMAN | BICD family-like cargo adapter 2 | Y2H |
| CYTSA_HUMAN | Cytospin-A | AP-MS |
| EPIPL_HUMAN | Epiplakin | AP-MS |
| ITBP2_HUMAN | Integrin beta-1-binding protein 2 | Y2H |
| LZTS2_HUMAN | Leucine zipper putative tumor suppressor 2 | Y2H |
| LMO7_HUMAN | LIM domain only protein 7 | AP-MS |
| ML12B_HUMAN | Myosin regulatory light chain 12B | AP-MS |
| MYL9_HUMAN | Myosin regulatory light polypeptide 9 | AP-MS |
| MYL6B_HUMAN | Myosin light chain 6B 208 | AP-MS |
| MYL6_HUMAN | Myosin light polypeptide 6 | AP-MS |
| PARP1_HUMAN | Poly(ADP-ribose) polymerase 1 | AP-MS |
| STON2_HUMAN | Stonin 2905 | AP-MS;Y2H |

Note that partners associated with ‘S active’ and ‘S active RBD1’ were excluded from the analysis because the paper by (Zhou *et al*., 2022) provided insufficient information regarding the precise nature of these two terms in both the main text and the Materials and Methods section. Including these interactions would have resulted in an expanded set of 25 proteins (adding ZNF274, ZDHHC5, and ELF4).

### Protein Data Bank entry 6XE9

Structure 6XE9 (https://www.rcsb.org/structure/6XE9) represents the heavy chain of Myosin II complex (Yang *et al*., 2020) of *Meleagris gallopavo* (wild turkey). Its UniProt accession number is G1N5L2 and its sequence length is 1979 amino acids. This protein, also referred to as “class II myosin heavy chain,” is a part of the “myosin II complex” which enables muscle cells to contract and facilitates movement and shape changes in non-muscle cells. This complex contains two class II myosin heavy chains, two myosin essential light chains, and two myosin regulatory light chains. These chains assemble to form an alpha-helical, coiled-coil tail.

G1N5L2 is not fully modeled; the protein segments 1-23, 205-210, 635-655, 955-1396, and 1676-1979 are missing. As a consequence:

- Protein segment 24-204 aligns with structure segment 1-181.
- Protein segment 211-634 aligns with structure segment 182-605.
- Protein segment 656-954 aligns with structure segment 606-904.
- Protein segment 1397-1675 aligns with structure segment 905-1183.

G1N5L2 is made of two InterPro domains: a Myosin Head Motor Domain (IPR001609, positions 78-790) and a Myosin Tail Domain (IPR002928, positions 854-1934). The alignments between the Myosin Head domain and the modeled part of this domain are the following:

- Protein segment 78-204 aligns with structure segment 55-181,
- Protein segment 211-634 aligns with structure segment 182-605,
- Protein segment 656-790 aligns with structure segment 606-740.

The alignments between the Myosin Tail domain and the modeled part of this domain are the following:

- Protein segment 854-954 aligns with structure segments 804-904.
- Protein segment 1397-1675 aligns with structure segments 905-1183.

Thus, the modeled Myosin Head covers structure segment 55-740, and the modeled Myosin Tail spans structure segment 804-1183.

### Phyre2: Protein structure prediction

Phyre2 (Protein Homology/AnalogY Recognition Engine), a free web-based service for protein structure prediction (Kelley *et al*., 2015), has been used for protein modeling (*i*.*e*., inferring the three-dimensional structure of a protein from its amino acid sequence).

### BackPhyre: Identification of proteins containing a given structural domain

The identification of potential Spike S interaction partners in the human proteome was carried out using the BackPhyre program (see <u>BackPhyre documentation</u>). BackPhyre takes a known 3D structure and searches a given proteome for proteins which harbor that specific structure (or a segment of it). I queried the human proteome (NCBI Reference Sequence database) using PDB structure 6XE9 (the heavy chain of myosin II complex). BackPhyre returned 999 protein entries with probabilities ranging from 95.8% to 100%. According to BackPhyre’s documentation, if a match probability exceeds 90%, one can generally be very confident that the protein adopts the overall fold shown and that the core of the protein is modelled at high accuracy (2-4Å rmsd from native, true structure).

From the original 999 entries, I excluded 195 sequences prefixed with “XP_,” indicating they were not curated RefSeq records. This left us with a set of 804 curated protein sequences (starting with “NP_”). I then grouped together the different isoforms of each protein-coding gene, resulting in 622 distinct protein clusters (see Supplemental file 2).

Given that I fed BackPhyre with the entire PDB structure 6XE9, and to the extent that the feature identified as common to 12 out of the 22 Spike S’ partners (see Results) concerns the C-terminal part of 6XE9, I performed a filtering step to preserve clusters that encompass at least one isoform possessing this characteristic. To achieve this, I identified the boundaries of the largest motif of 6XE9 common to all of my 12 previously referred proteins; I found 853-1067. This step narrowed down the set to 488 protein clusters. Finally, to represent each protein cluster uniquely, I selected a single isoform from those with multiple isoforms.

Note that BackPhyre’s human protein-coding gene set is based on the <u>NCBI36/hg18</u> genome assembly (released March 3, 2006), an older version than the current <u>GRCh38.p14</u> assembly (released February 3, 2022). Although incomplete, this release is sufficiently accurate, as only 5 of my set of 488 identified human proteins have no UniProt ID or correspond to an obsolete entry (see Supplemental file 3), though the annotations of the majority of the remaining 483 proteins were obviously subsequently updated. Consequently, my analysis remains sound, though BackPhyre’s protein set may not be fully representative of human coiled-coil proteins.

## Statistical analysis

To evaluate whether a given set of proteins of size N was statistically enriched for a specific feature F compared to its expected distribution within the human proteome, I performed a Fisher’s exact test (significance level = 0.05). Specifically, considering that a number n of proteins from the given set share a common feature F, I tested the null hypothesis that this set is a random sample of size N drawn from the human proteome. Defining N_H as the size of the human proteome and n_H as the number of human proteins possessing feature F, the structure of the contingency table and the corresponding formulas for the counts are as follows:

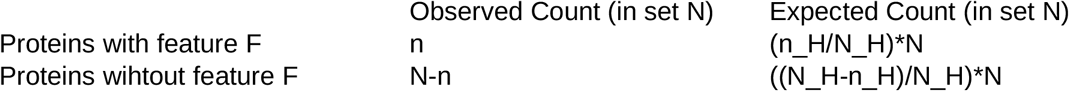

Fisher’s exact test was conducted on the website BiostatTGV (https://biostatgv.sentiweb.fr/?module=tests/fisher).

According to UniProt (UniProt Consortium, 2024), the total number of proteins in the human proteome is 82518 (query: (proteome:UP000005640), where UP000005640 is the UniProt ID of the human proteome).

I used UniProt Gene Ontology queries to define the feature F and to compute the number of human proteins with this feature. The human proteins with the feature ‘intermediate filament protein’ were identified using the query: (proteome:UP000005640) AND (go:0005882), where 0005882 is the GO term for ‘intermediate filament’. Similarly, human proteins with the feature ‘kinesin’ were identified using the query: (proteome:UP000005640) AND (go:0005871), where 0005871 is the GO term for ‘kinesin complex’.

For the feature ‘actin-binding protein’, I relied on a collection of 418 actin-associated proteins (AAPs) whose direct interactions with actin have been experimentally verified (Gao & Nakamura, 2022).

## Results

### Spike S partners are enriched in cytoskeleton-associated proteins, particularly actin-binding proteins, which are statistically overrepresented compared to their distribution in the human proteome

Among the 22 Spike S partners (see Table 2 for a summary), 10 are annotated as “Actin Binding Proteins” according to a study which gathered a collection of 418 actin-associated proteins (AAPs) whose direct interactions with actin have been experimentally verified (Gao & Nakamura, 2022). These 10 proteins play various roles in actin dynamics (see UniProt annotations (UniProt Consortium, 2025), (Gao & Nakamura, 2022) and (Cooper, 2000)):

**Table 2.**
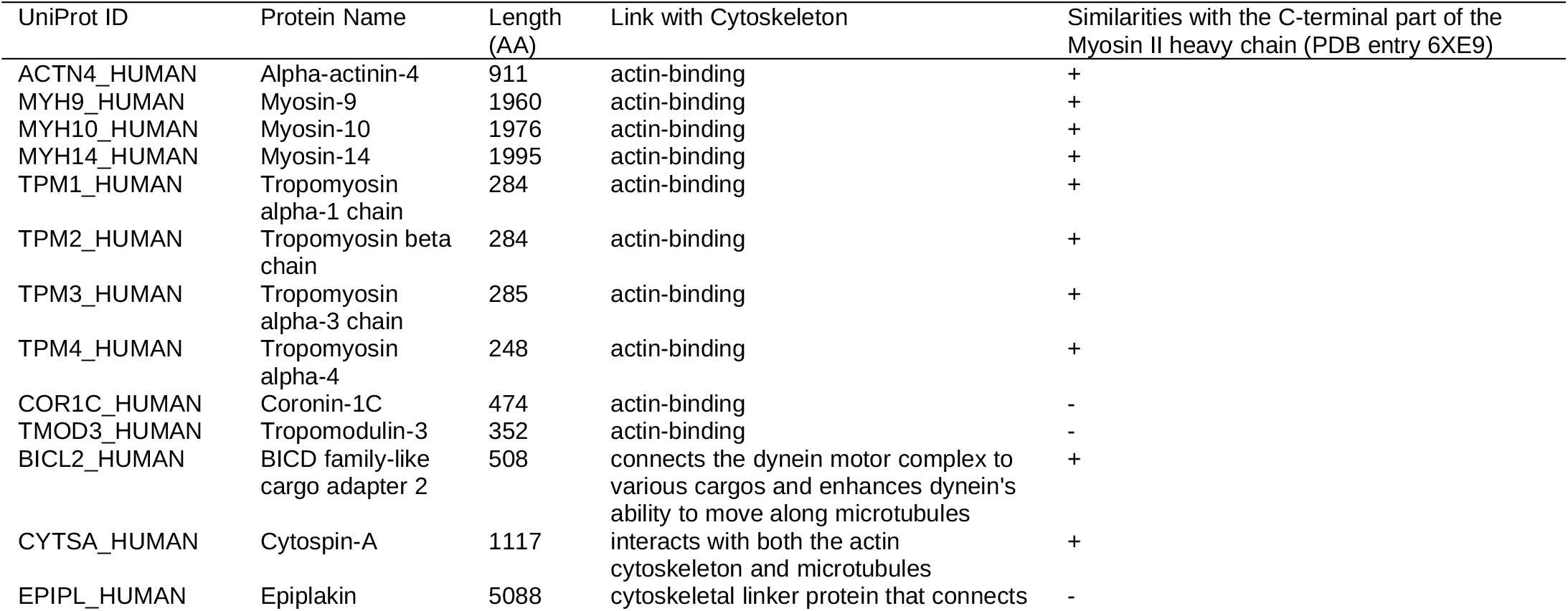
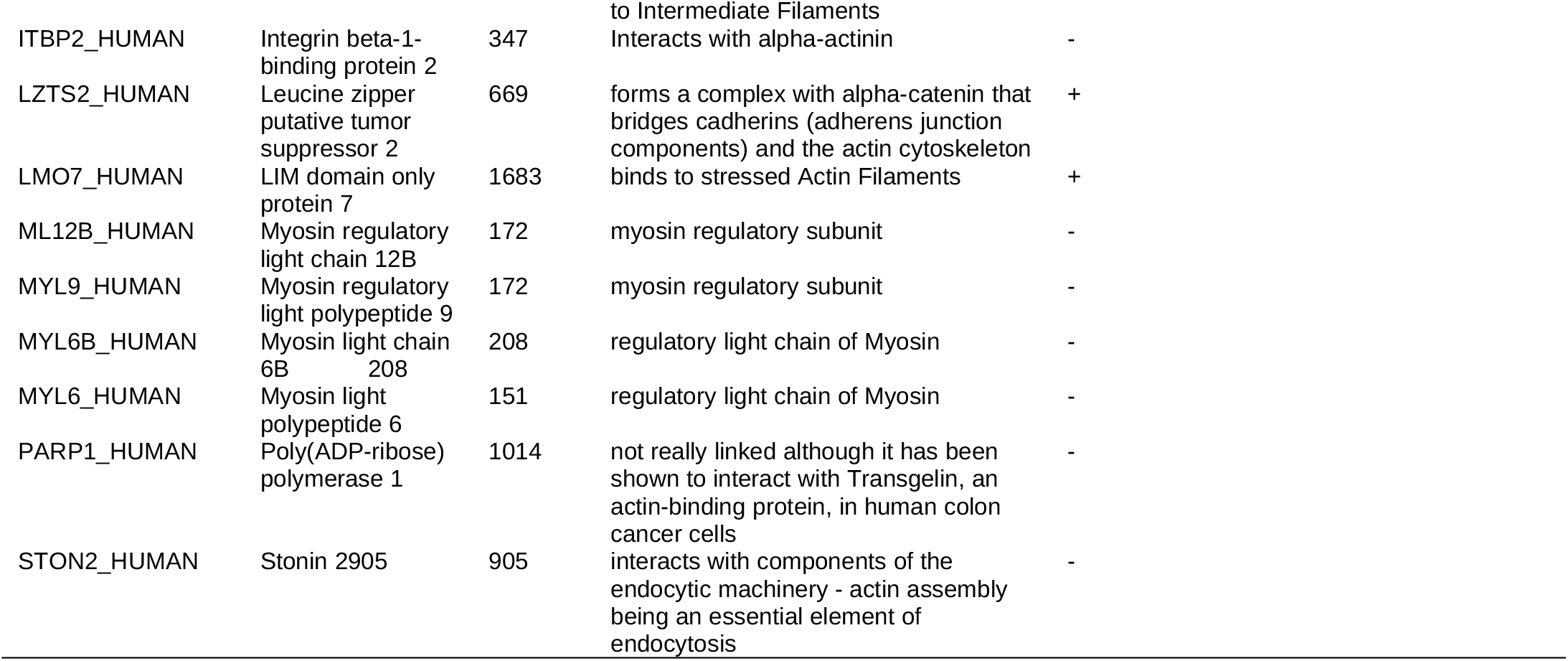
Each line represents a Spike S partner. The ‘UniProtID’ column identifies the protein using its unique identifier in the UniProt database. The ‘Protein Name’ column indicates the name of the protein. The ‘Length (AA)’ column indicates the total length of the protein sequence (in amino acids). The ‘Link with Cytoskeleton’ column indicates whether a documented link exists between the considered Spike S partner and the cytoskeleton. The ‘Similarities with the C-terminal part of Myosin II heavy chain (PDB entry 6XE9)’ column indicates whether the considered Spike S partner shares similarities (+) or does not share similarities (-) with the C-terminal tail of the Myosin II heavy chain (this latter column refers to subsection ‘A majority of Spike S partners share similarities with the C-terminal myosin II heavy chain domain’).

- “Alpha-actinin-4” (UniProt ID ACTN4_HUMAN) is a component of the actin cytoskeleton.
- “Myosin-9” (UniProt ID MYH9_HUMAN), “Myosin-10” (MYH10_HUMAN), and “Myosin-14” (MYH14_HUMAN) are cellular myosin.
- “Tropomyosin alpha-1 chain” (UniProt ID TPM1_HUMAN), “Tropomyosin beta chain” (TPM2_HUMAN), “Tropomyosin alpha-3 chain” (TPM3_HUMAN), and “Tropomyosin alpha-4 chain” (TPM4_HUMAN) are involved in stabilizing actin filaments (e.g., protecting F-actin from severing).
- “Coronin-1C” (UniProt ID COR1C_HUMAN) and “Tropomodulin-3” (TMOD3_HUMAN) are involved in stabilizing, bundling, and capping actin, respectively.

In the ‘Statistical Analysis’ section, I demonstrate a statistically significant enrichment of actin-binding proteins among the set of Spike S protein interactors compared to their expected distribution within the human proteome.

The remaining 12 Spike S partners interact directly or indirectly with the cytoskeleton, an intricate network of filaments within cells comprising actin filaments (AFs), microtubules (MTs), and intermediate filaments (IFs) (Cooper, 2000):

- “BICD family-like cargo adapter 2” (UniProt ID BICL2_HUMAN) belongs to the BICD family-like. “BICD family-like cargo adapter 1” (BICL1_HUMAN), a member of this family, is predicted to connect the dynein motor complex to various cargos and enhance dynein’s ability to move along microtubules (MTs) for extended distances without falling off (see UniProt annotations).
- “Cytospin-A” (UniProt ID CYTSA_HUMAN) is documented as a “cross-linking” protein that functionally interacts with both the actin cytoskeleton and MTs (Saadi *et al*., 2011).
- “Epiplakin” (UniProt ID EPIPL_HUMAN) is a cytoskeletal linker protein that connects to IFs and controls their reorganization in response to stress ((Jang *et al*., 2005), (Shimada *et al*., 2013), (Kokado *et al*., 2016)).
- “Integrin beta-1-binding protein 2” (UniProt ID ITBP2_HUMAN), also referred as Melusin, is a muscle-specific interactor for the beta(1) integrin’s cytoplasmic domain, which interacts with the actin-binding protein alpha-actinin (Otey *et al*., 1990)
- “Leucine zipper putative tumor suppressor 2” (UniProt ID LZTS2_HUMAN) is a beta-catenin-interacting protein that modulates beta-catenin signaling and localization (Thyssen *et al*., 2006). Beta-catenin, a crucial regulator of cell-cell adhesion, forms a complex with alpha-catenin that bridges cadherins (adherens junction components) and the actin cytoskeleton (Kobielak & Elaine Fuchs, 2004).
- “LIM domain only protein 7” (UniProt ID LMO7_HUMAN) belongs to a group of proteins that bind to stressed actin filaments (Winkelman *et al*., 2020).
- “Myosin regulatory light chain 12B” (UniProt ID ML12B_HUMAN) and “Myosin regulatory light polypeptide 9” (MYL9_HUMAN) are part of the Myosin II complex, as are “myosin light chain
- 6B” (MYL6B_HUMAN) and “myosin light polypeptide 6” (MYL6_HUMAN). These light chains do not bind to actin but interact with the myosin heavy chain, which allow Myosin II complex to bind to actin.
- “Poly(ADP-ribose) polymerase 1” (UniProt ID PARP1_HUMAN), which is mainly and well documented to play a key role in DNA repair, also interacts with Transgelin, an actin-binding protein, in human colon cancer cells (Lew *et al*., 2020). Note that several studies underscore the functional importance of lamins, actin, myosin, spectrin, and the linker of nucleoskeleton and cytoskeleton (LINC) complex in DNA repair (Lambert, 2019). Finally note that while PARP1 is involved in multiple antiviral mechanisms, several viruses can inhibit these antiviral functions and recruit and utilize PARP1 for their own benefit (Sobotka & Tempera, 2024).
- “Stonin 2” is an adaptor protein that interacts with components of the endocytic machinery (Martina *et al*., 2020), which are involved in regulating coat assembly, vesicle budding, and signal transduction pathways along with the actin cytoskeleton (Chakrabarti *et al*., 2021).

A Fisher’s exact test was performed to assess the enrichment of ‘actin-binding’ proteins within the set of 22 Spike S partners (10 of which are actin-binding proteins). With a calculated Fisher’s exact p-value of 0.000521 (significance level = 0.05), the null hypothesis is rejected, indicating that actin-binding proteins are significantly overrepresented in this set relative to the entire human proteome.

In conclusion, the set of 22 Spike S partners is enriched in “cytoskeleton-associated proteins”, particularly in actin-binding proteins which are statistically over-represented.

### A majority of Spike S partners share similarities with the C-terminal Myosin II heavy chain domain

The observation that a protein interacts with a set of proteins often leads to the speculation that these proteins share a common domain mediating the identified interactions. This is why I investigated whether a majority of the 22 Spike S protein partners would share a common domain. Using InterPro domain annotations, I found no such domain. Therefore, I decided to use Phyre2, a protein 3D structure prediction method (Kelley *et al*., 2015), to address this question.

I ran Phyre2 on each of the 22 Spike S partners (see Supplemental file 1). Interestingly, a careful manual examination of the results revealed that 12 out of the 22 Spike S partners shared a high-confidence similarities with the C-terminal part of the PDB entry 6XE9, *i*.*e*., the myosin II heavy chain of the myosin II complex of *Meleagris gallopavo* (wild turkey) (see Materials and Methods section ‘Protein Data Bank entry 6XE9’ for a detailed presentation of 6XE9). Its UniProt accession number is G1N5L2 and its sequence length is 1979 amino acids (see (Yang *et al*., 2020)). It is made of two InterPro domains: a Myosin Head Motor Domain (IPR001609, positions 78-790) and a Myosin Tail Domain (IPR002928, positions 854-1934). As highlighted in subsection ‘Protein Data Bank entry 6XE9’, 6XE9 is not fully modeled. Specifically, the modeled Myosin Head covers structure segments 55-740, and the modeled Myosin Tail spans structure segments 804-1183.

The 12 Spike S partners that share similarities with the C terminal part of the PDB entry 6XE9 are listed in the Table 3 below. For each of these latter, this table summarizes some protein characteristics (UniProt ID, name, length) as well as its alignment with PDB entry 6XE9 and Phyre2’s confidence score.

**Table 3.**
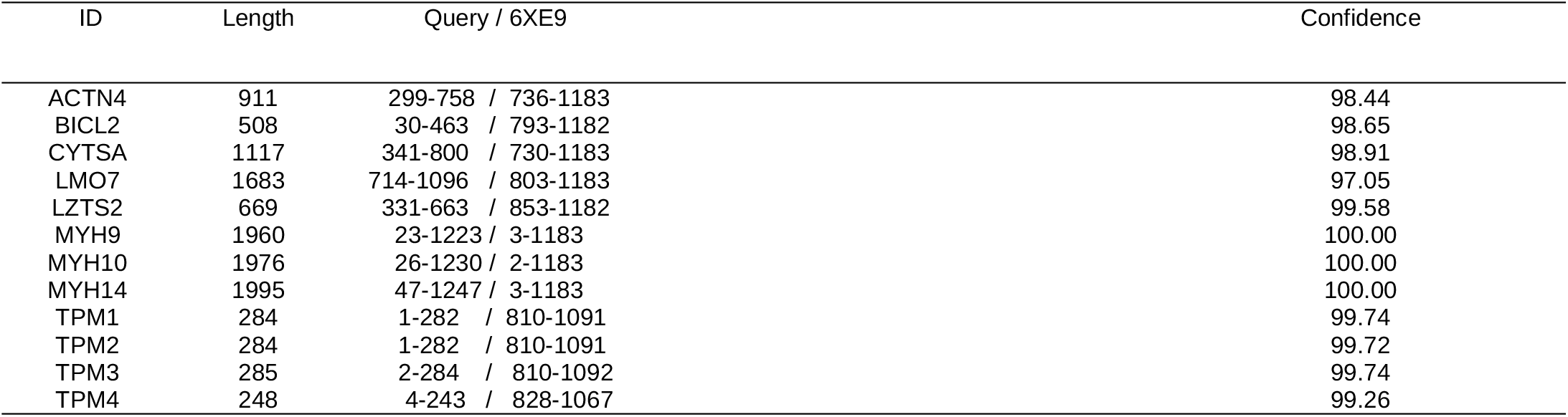
Each line represents a Spike S partner exhibiting similarities with the C-terminal part of PDB entry 6XE9 (Myosin II heavy chain). The ‘ID’ column provides the protein’s UniProt identifier. The ‘Length’ column indicates the protein’s length (in amino acids). The ‘Query/6XE9’ column shows the aligned amino acid positions between the query protein and the reference structure (6XE9), in the format start-end (query)/start-end (6XE9). The ‘Confidence’ column displays the Phyre2 confidence score for the similarity, indicating the probability of accurate alignment.

The results reveal a clear fit between protein structure 6XE9 and myosin heavy chains MYH9, 10, and 14, which is consistent as these proteins belong to the same family. The tropomyosins (TPM1, 2, 3, and 4) exhibit a significant fit with the tail of 6XE9, leaving only a minor portion of their sequences not covered. A substantial portion of the actin sequence ACTN4 also aligns with 6XE9 tail. Finally, the similarity shared between 6XE9 and BICL2 (resp., CYSTA, LMO7, and LZTS2) is localized in the tail region of 6XE9. In conclusion, these 12 proteins share similarities with the Myosin tail domain of 6XE9.

### The set of human proteins identified by BackPhyre is enriched in cytoskeletal-associated proteins, with actin-binding proteins, intermediate filament proteins, and kinesins all being statistically overrepresented compared to their distribution in the human proteome

To identify potential human interactors/partners of Spike S, I employed BackPhyre which, given a 3D structure, determines if this structure (or part of this structure) is present in proteins of a genome/proteome of interest. I queried the human proteome (NCBI Reference Sequence database) using PDB structure 6XE9 (the heavy chain of myosin II complex), see Materials and Methods, ‘BackPhyre: Identification of proteins containing a given structural domain’ section.

BackPhyre returned a set of 999 high-confidence hits (probability greater than 0.97). Following a cleaning step (excluding RefSeq entries annotated as ‘XP’ and grouping together isoforms), I obtained a set of 488 human proteins, which I then clustered into 7 major groups.

#### 1. / ‘Link To Actin’ cluster

I relied on a list of manually curated Actin-Associated Proteins (AAPs) (Gao & Nakamura, 2022), which contain Actin-Binding Proteins (ABPs), to identify the actin-binding proteins of my set. This resulted in a cluster that includes proteins known to interact with actin, such as actin, myosin, and tropomyosin. Additionally, I also included proteins functionally linked to actin but not explicitly annotated as actin-binding proteins. The ‘Link To Actin’ cluster contains 67 proteins, among which 56 are ABPs; I demonstrate (see ‘Statistical analysis’) that my set of 488 proteins is statistically enriched with ABPs compared to their distribution in the human proteome.

#### 3. / ‘Link to Intermediate Filaments’ cluster

I relied on UniProt annotations to identify Intermediate Filament (IF) proteins within the human proteome. (query: “(proteome:UP000005640) AND (go:0005882) where ‘0005882’ refers to ‘intermediate filament’ in Gene Ontology). ‘Link to Intermediate Filaments’ cluster groups together IFs, such as keratins, vimentin, neurofilament proteins, etc., along with proteins linked to intermediate filaments. ‘Link to Intermediate Filaments’ contains 71 proteins among which 68 are IFs. I show in ‘Statistical analysis’ that my set of 488 proteins is statistically enriched with IFs compared to their distribution in the human proteome.

#### 3. / ‘Link To Kinesin and Dynein’ cluster

My analysis of my set of 488 proteins revealed an absence of tubulins, the building blocks of microtubules. Despite this, I identified several kinesin and dynein motor proteins, known for their interaction with microtubules to facilitate intracellular transport of various cargo like vesicles, organelles, and chromosomes. I additionally put four ‘Tektins’ in this cluster. According to UniProt annotations, these latter are microtubule inner proteins (MIPs) that are components of the dynein-decorated doublet microtubules (DMTs) in cilia and flagellar axonemes. The ‘Link To Kinesin and Dynein’ cluster contains 46 proteins, among which 24 are kinesins (query: “(proteome:UP000005640) AND (go:0005871)” where ‘0005871’ refers to ‘kinesin complex’ in Gene Ontology). I prove in section ‘Statistical analysis’ that my set of 488 proteins is statistically enriched with kinesins compared to their distribution in the human proteome.

#### 4. / ‘Link To Microtubule’ cluster

The identification of microtubule motor proteins (kinesins and dyneins, as shown in the cluster above) prompted us to search for microtubule-related proteins in my set of 488 proteins. Microtubules, anchored at the centrosome (the MTOC), extend towards the cell periphery and play a vital role in various cellular processes, e.g., during mitosis, they reorganize into the mitotic spindle, ensuring proper chromosome segregation. Additionally, cilia and flagella are microtubule-based structures that contribute to cellular functions. Not surprisingly, my analysis identified many proteins associated with these structures (centrosome, mitotic spindle, cilia, and flagella). These proteins were subsequently grouped together into 5 different subclusters, namely, ‘Link to MT’, Link To Centrosome’, ‘Link To Cilia’, ‘Link To “Sperm Flagellar”‘, and ‘Link To Spindle’. The ‘Link To Microtubule’ cluster contains 104 proteins.

#### 5. / ‘Link To Trafficking’ cluster

Functional analysis of the remaining proteins allowed us to identify an enrichment in proteins associated with transport and trafficking pathways. The ‘Link To Trafficking’ cluster contains 51 proteins, among which are several members of the BICD family. Members of this family are known to link the dynein motor complex to various cargos.

#### 5. ‘Various’ cluster

I next identified several clusters with a moderate number of proteins, associated for instance with ‘defense’ or ‘LINC (Linker of Nucleoskeleton and Cytoskeleton)’, this latter grouping 3 proteins involved in the connection between the nuclear lamina and the cytoskeleton. The ‘Various’ cluster contains 106 proteins.

#### 7. ‘Lacking UniProt Summary’ cluster

This cluster contains 43 proteins lacking a UniProt ‘Function’ field, which was one of my main source of information for performing my clustering.

To evaluate whether the set of 488 human proteins identified by BackPhyre was statistically enriched for specific functional categories, I performed Fisher’s exact tests. The analysis demonstrated that three key cytoskeletal protein classes, namely actin-binding proteins (grouped in the ‘Link To Actin’ cluster, p-value = 1.307E-14), intermediate filament proteins (grouped in the ‘Link to Intermediate Filaments’ cluster, p-value = 3.935E-19), and kinesins (grouped in the ‘Link To Kinesin and Dynein’ cluster, p-value = 8.921E-8), are significantly overrepresented (significance level 0.05) (see Materials and Methods, Statistical analysis section).

## Discussion

To gain deeper insights into SARS-CoV-2 infection dynamics, I re-examined the most comprehensive SARS-CoV-2–human protein interactome (Zhou *et al*., 2022), which leverages both Y2H (Rajagopala *et al*., 2015) and AP-MS (Causier *et al*., 2004) methods. I focused on the Spike S protein, the viral entry protein that binds to the host cell receptor ACE2.

While Y2H and AP-MS have limitations (see Introduction), they generally produce high-quality data. AP-MS identifies protein complexes while Y2H reveals binary interactions, providing complementary insights. Nevertheless, initially, I decided to treat Y2H and AP-MS equally as two probes of interaction. This decision was primarily driven by the observation that Human Myosin-9, identified as a Spike S partner through AP-MS in the interactome I utilized (Zhou *et al*., 2022), had previously been demonstrated to be in close proximity to the SARS-CoV-2 Spike S protein (using an APEX2 proximity-labeling technique) and subsequently showed to directly interact with it (Chen *et al*., 2021).

Zhou and colleagues’ interactome identified 22 Spike S interactors/partners (see Materials and Methods section ‘Interactome data’ and Table 1). Analysis of this set of 22 partners revealed a predominance of cytoskeleton-associated proteins, with a statistically significant overrepresentation of actin-binding proteins (10 out of 22) compared to their distribution in the human proteome (see Results, ‘Spike S partners are enriched in cytoskeleton-associated proteins, particularly actin-binding proteins, which are statistically overrepresented compared to their distribution in the human proteome’ section, and Table 2).

To further investigate the interactome results, I hypothesized that the human Spike S partners might possess a common domain mediating these interactions. Unable to identify such domains using InterPro annotations (Paysan-Lafosse *et al*., 2022), I used Phyre2 (Kelley *et al*., 2015) to predict the structures of the 22 Spike S partners. I then observed that 12 of these 22 structures exhibited high-confidence similarities with the tail region/domain of the Myosin II heavy chain (PDB entry 6XE9, see Results,’A majority of Spike S partners share similarities with the C-terminal Myosin II heavy chain domain’ section, and Table 3). Among these 12 proteins, 8 are actin-binding proteins, including human Myosin-9, which was previously shown to interact with the Spike S protein via its C-terminal domain (as mentioned in the previously).

My analysis reveals several key findings: 1) Actin-binding proteins are significantly overrepresented among the 22 Spike S partners (10 among 22), 2) Eight of these actin-binding proteins exhibited structural similarities to the C-terminal region of Myosin II heavy chain, 3) Myosin-9, a cellular myosin and one of these eight proteins, was previously shown to interact with the Spike S protein through its C-terminal domain (Chen *et al*., 2021). Importantly, this C-terminal domain of Myosin-9 is homologous to the C-terminal region of Myosin II heavy chain which is a well-documented coiled-coil region (Rahmani *et al*., 2021). These findings led us to hypothesize that domains homologous to the C-terminal region of Myosin II heavy chain may play a crucial role in mediating interactions with the Spike S protein.

While proteins lacking this type of domain might still interact with the Spike S protein through other, yet unidentified motifs or domains, my hypothesis provides the most parsimonious explanation for the observed data. Specifically, this hypothesis clarifies why Affinity Purification-Mass Spectrometry (AP-MS), which detects protein complexes rather than individual protein-protein interactions, might identify Spike S partners that don’t exhibit similarities to the C-terminal region of Myosin II heavy chain. This is exemplified by Myosin II light chains. Myosin II light and heavy chains are integral components of the Myosin II complex. Given that the Myosin II complex binds to actin through its heavy chains, the identification of light chains by AP-MS likely reflects their association with the heavy chains within the complex, rather than indicating a direct interaction between the Spike S protein and the light chains themselves.

To identify potential human Spike S protein partners, I decided to search for human proteins sharing similarities with the C-terminal part of Myosin II heavy chain. To achieve this, I used BackPhyre (‘Phyre2 in reverse’). Given a 3D protein structure, this tool searches for homologous sequences across a broad range of genomes/proteomes. I ran BackPhyre with the PDB entry 6XE9 (corresponding to the Myosin II heavy chain) to explore the human proteome (see Materials and methods, section BackPhyre: Identification of proteins containing a given structural domain).

BackPhyre returned a set of 999 high-confidence hits (probability >= 0.97). Following a cleaning step (excluding RefSeq entries annotated as ‘XP’ and grouping together isoforms), I obtained a set of 488 human proteins, which I then clustered into 7 major groups. I next demonstrated that three major classes of cytoskeletal proteins are statistically overrepresented in this set compared to their distribution in the human proteome: Actin-binding proteins (grouped in the ‘Link To Actin’ cluster), Intermediate filament proteins (grouped in the ‘Link to Intermediate Filaments’ cluster), and Kinesins (grouped in the ‘Link To Kinesin and Dynein’ cluster) (see Results, ‘The set of human proteins identified by BackPhyre is enriched in cytoskeletal-associated proteins, with actin-binding proteins, intermediate filament proteins, and kinesins all being statistically overrepresented compared to their distribution in the human proteome’ section).

These findings are consistent with existing research emphasizing the importance of the cytoskeleton in viral infections (Kloc *et al*., 2022; Wen *et al*., 2022). Hence, the ‘Link To Actin’ cluster contains the non-muscle myosin heavy chain IIA (MYH9), which has been shown to interact with Spike S and to facilitate SARS-CoV-2 infection in human pulmonary cells (Chen *et al*., 2021). The ‘Link to Intermediate Filaments’ cluster contains vimentin, which has also been documented to interact with Spike S and to facilitate entry into human endothelial cells (Amraei *et al*., 2022). The ‘Link To Kinesin and Dynein’ cluster contains kinesin-1, a protein documented to be hijacked by several viruses for transport along microtubules (to reach their destinations in the highly crowded cellular environment). For instance, the intercellular transport of Porcine Epidemic Diarrhea Virus (PEDV), which is a coronavirus, is driven along microtubules by both dynein and kinesin-1 (Hou *et al*., 2021). Furthermore, cytoplasmic dynein and kinesin-1 have both been found to bind Adenovirus (AdV) through direct interactions with the capsid proteins (Scherer *et al*., 2020). Although, to the best of my knowledge, no study has yet reported that SARS-CoV-2 hijacks kinesin-1 via an interaction with Spike S, exploring this possibility represents an interesting avenue for future research.

In conclusion, my work provides *in silico* evidence suggesting that SARS-CoV-2 Spike S might target proteins harboring regions homologous to the C-terminal region of Myosin II heavy chain, which is a coiled-coil domain. These regions, as shown here, are particularly prevalent in cytoskeletal and cytoskeleton-associated proteins. These regions, as shown here, are particularly prevalent in cytoskeletal and cytoskeleton-associated proteins (for example, myosins, intermediate filament proteins, and kinesins; see also Simm *et al*., 2021) and are documented to be hijacked by several viruses. However, since this work is based primarily on a large-scale protein-protein interaction dataset, which may contain false positives or interactions irrelevant in physiological contexts (e.g., non-co-localization), direct biological extrapolations require caution. Further biochemical and cell biological investigations are necessary to study the interaction between Spike S and proteins containing coiled-coil domains, and the involvement of such domains in these interactions.

## Acknowledgements

I warmly thank Joël Pothier (ABI, ISYEB) for his careful review of a draft of this manuscript. I am also grateful to the two referees of my paper, whose comments and suggestions significantly improved the quality of the final submission.

## Supplementary information availability

Supplemental Files 1, 2, and 3 are available online (https://doi.org/10.5281/zenodo.14718017).

---

**Figure.**
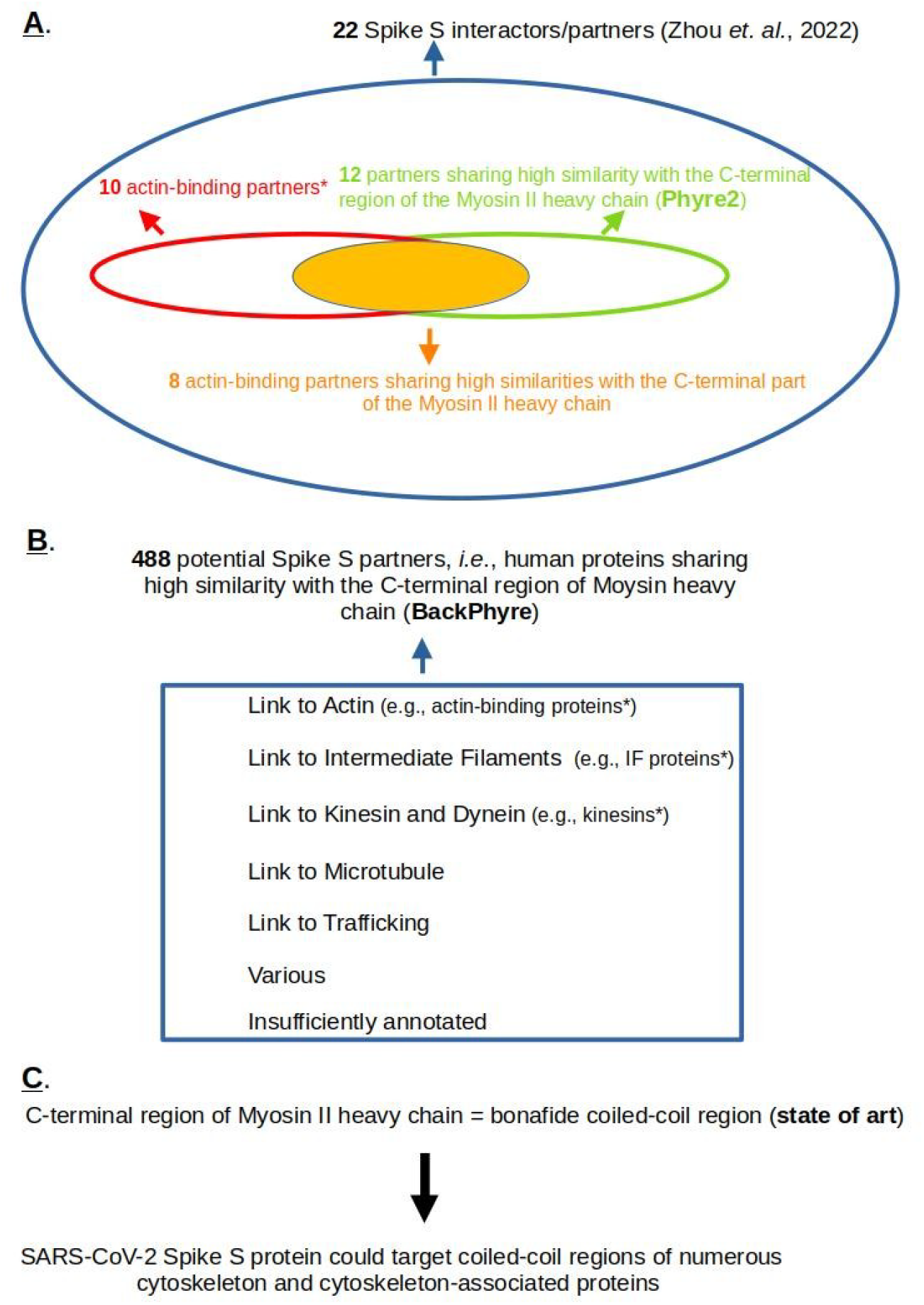
A. Focusing on a human ‘Spike S interactors/partners’ set of 22 proteins (Zhou *et al*., 2022), I show that actin-binding proteins are statistically overrepresented (10 among 22) compared to their distribution in the human proteome. Next, using Phyre2 (Kelley *et al*., 2015), a bioinformatics tool for protein structure prediction, I identify that 12 out of 22 Spike S partners contain a structural domain sharing similarities with the C-terminal part of the Myosin II heavy chain. B./ Using BackPhyre (Kelley *et al*., 2015), a tool that identifies proteins harboring a given structural domain, I identified a set of 488 human proteins that, with high probability, share similarites with the C-terminal part of the Myosin II heavy chain. This set of proteins is particularly abundant in human cytoskeletal and associated proteins (statistically overrepresented in actin-binding proteins, Intermediate Filament proteins, and kinesins), as well as in proteins involved in microtubule dynamics. C. Given that the C-terminal domain of the Myosin II heavy chain is documented as a coiled-coil region, I infer that Spike S might target proteins containing such regions. * means statistically overrepresented.

